# Ras Suppresses TXNIP Expression by Restricting Ribosome Translocation

**DOI:** 10.1101/299354

**Authors:** Zhizhou Ye, Donald E. Ayer

## Abstract

Oncogenic Ras upregulates aerobic glycolysis to meet the bioenergetic and biosynthetic demands of rapidly growing cells. In contrast, Thioredoxin interacting protein (TXNIP) is a potent inhibitor of glucose uptake and is frequently downregulated in human cancers. Our lab previously discovered that Ras activation suppresses TXNIP transcription and translation. In this report, we developed a system to study how Ras affects TXNIP translation in the absence of transcriptional affects. We show that whereas Ras drives a global increase in protein translation, it suppresses TXNIP protein synthesis by reducing the rate at which ribosomes transit the coding region of TXNIP mRNA. To investigate the underlying mechanism(s), we randomized or optimized the codons in the TXNIP message without altering the TXNIP primary amino acid sequence. Translation from these mRNA variants is still repressed by Ras, intimating that mRNA secondary structure, miRNAs, RNA binding proteins, or codon usage do not contribute to the blockade of TXNIP synthesis. Rather, we show that the N-terminus of the growing TXNIP polypeptide is the target for Ras-dependent translational repression. Our work demonstrates how Ras suppresses TXNIP translation elongation in the face of a global upregulation of protein synthesis and provides new insight into Ras-dependent metabolic reprogramming.

## INTRODUCTION

Activating mutations in the small Ras GTPases (K-Ras, H-Ras, N-Ras) are among the most common alterations in human cancer. The oncogenic mutations render the Ras proteins constitutively active, which drives uncontrolled proliferation through the activation of the downstream signaling pathways, such as the MAPK and PI3K pathways (1, 2). Ras activation also rewires metabolism to accommodate the increased anabolic demands of rapidly growing and dividing cells. For example, Ras stimulates glucose uptake and aerobic glycolysis for ATP generation, diverts glycolytic intermediates into biosynthetic pathways, and upregulates glutaminolysis to fuel central carbon metabolism (3-5). The glycolytic switch conferred by Ras activation has been classically ascribed to its activation of c-Myc and HIF-1α, which directly drive the expression of glucose transporters and glycolytic enzymes (6-8). More recent transcriptional analysis showed that oncogenic Ras activates the expression of genes involved in a spectrum of anabolic pathways, including the hexosamine, ribose and pyrimidine biosynthetic pathways (9). Each of these biosynthetic pathways is fueled by glucose-derived carbons. However, in the absence of glucose availability, flux through these biosynthetic pathways is limited. Thus, nutrient use must be coupled with nutrient availability to sustain the rapid growth and division of transformed cells.

Thioredoxin interacting protein (TXNIP) is a critical negative regulator of cellular glucose uptake. It inhibits glucose uptake by removing glucose transporters from the cell surface (10, 11). Consequently, TXNIP loss is sufficient to drive glucose uptake and aerobic glycolysis (12-15). Further, TXNIP can also promote oxidation of non-glucose fuels (16, 17). Therefore, low TXNIP levels support the use of glucose as a fuel, whereas high TXNIP levels support the use of non-glucose fuels. In addition to this function in fuel choice, TXNIP has a number of additional anti-proliferative activities. For example, it can drive apoptosis by activating Ask1 and it can drive cell cycle arrest by stabilizing p27^kip1^ (18, 19). Given these assorted functions, it is not surprising that TXNIP functions as a tumor suppressor and is downregulated in a variety of human cancers (20-22).

Transcription of the TXNIP gene is highly, if not entirely, dependent on the MondoA transcription factor (20, 23). MondoA is a member of the extended Myc network and its activity is driven by high glucose (24). Thus, the MondoA-TXNIP axis constitutes a negative feedback loop that regulates glucose homeostasis (15, 25, 26). Our previous work established that the MondoA-TXNIP axis is downregulated by PI3K, mTOR or Myc activation, which contributes to their well characterized activities in driving aerobic glycolysis (20, 27, 28). TXNIP is also subject to post-translational regulation in response to signaling pathways. For example, under energy depletion, AMPK phosphorylates TXNIP leading to its degradation, permitting increased glucose uptake to restore energy homeostasis (10). The general model that emerges from these studies is that pro-growth signals downregulate TXNIP, which subsequently supports aerobic glycolysis and, presumably, anabolic reactions.

Our lab has previously shown that Ras activation downregulates TXNIP mRNA and protein expression (27). We propose that TXNIP repression provides an additional route to increase glucose uptake and utilization in response to Ras activation. In this report, we investigate how Ras inhibits TXNIP expression, with a focus on post-transcriptional regulation. We show that in spite of driving increased global translation, Ras suppresses translation elongation of TXNIP mRNA by targeting the N-terminus of nascent TXNIP polypeptide chain as it exits the ribosome exit tunnel.

## RESULTS

### Ras^G12V^ inhibits TXNIP expression

Our previous data suggest that acute Ras activation in immortalized human fibroblasts suppresses TXNIP transcription and translation (27). Consistent with Ras blocking TXNIP transcription, we observed a negative correlation between an H-Ras gene signature and TXNIP expression in breast cancer, lung adenocarcinoma and pancreatic adenocarcinoma patient samples using publicly available datasets (Fig.1A). Experimentally, expression of activated H-Ras (Ras^G12V^) in murine embryonic fibroblasts (MEFs) completely abolished TXNIP mRNA and protein expression (Fig.1B and C). The complete suppression of TXNIP transcription prevented us from investigating whether Ras^G12V^ regulates TXNIP post-transcriptionally.

**FIG 1.**
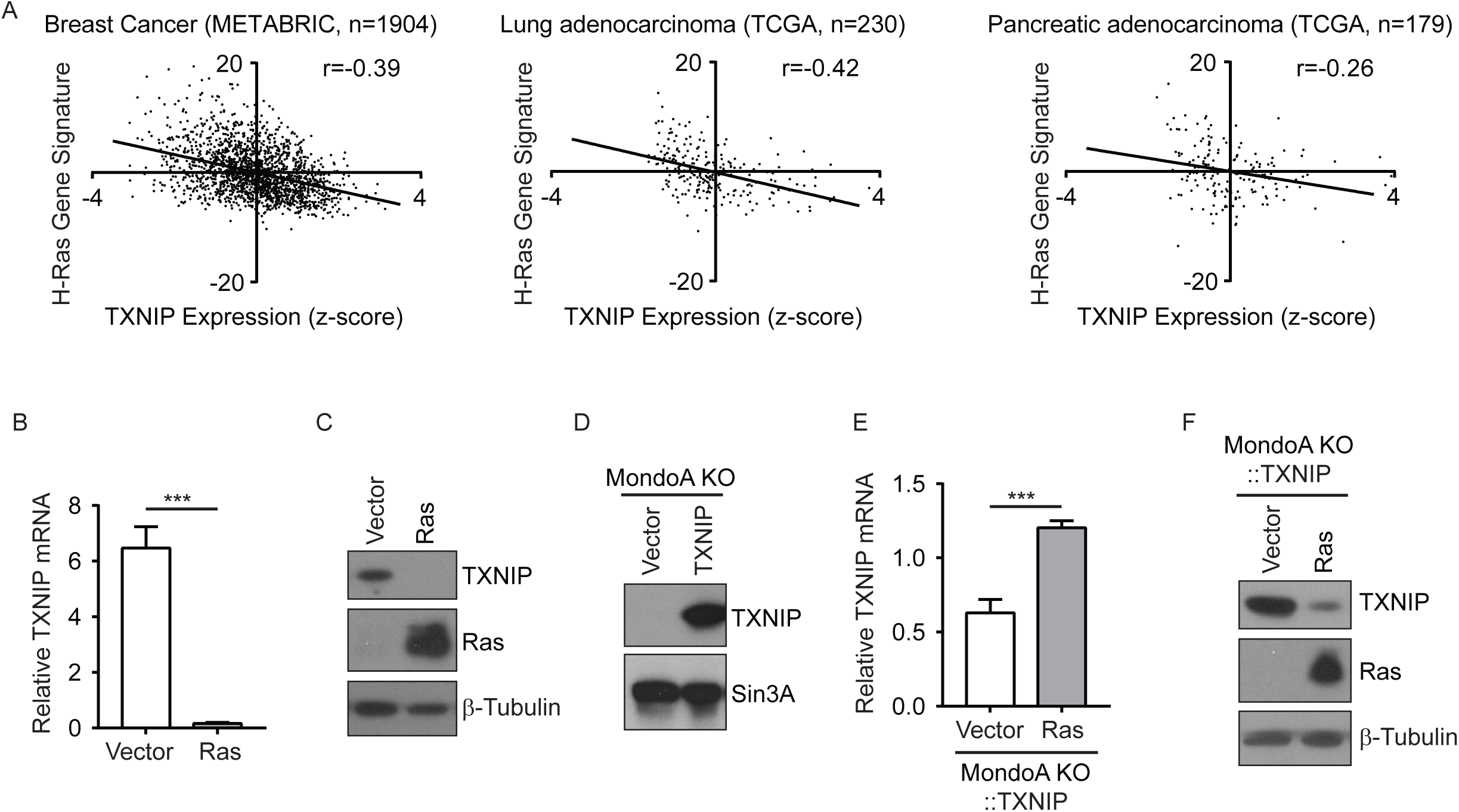
Ras^G12V^ inhibits TXNIP expression. **(A)** Scatter plots showing the relationship between a H-Ras gene signature and TXNIP mRNA expression level. Each point represents a single tumor sample from the indicated cancer datasets, with sample size denoted. The correlation coefficient Pearson’s r is shown for each dataset. **(B-C)** RT-qPCR (B) and western blotting (C) were used to determine relative TXNIP mRNA level (normalized to β-actin) and levels of the indicated proteins, respectively, in wildtype murine embryonic fibroblasts (MEFs) stably transduced with pBabePuro (Vector) or pBabePuro-H-Ras^G12V^ (Ras). **(D)** Western blotting was used to determine the levels of TXNIP and Sin3A proteins in MondoA knockout (MondoA KO) MEFs stably transduced with pWzlBlast (Vector) or pWzlBlast-TXNIP CDS (TXNIP). **(E and F)** MondoA KO::TXNIP MEFs were stably transduced with pBabePuro (Vector) or pBabePuro-H-Ras^G12V^ (Ras) as indicated. RT-qPCR (E) and western blotting (F) were used to determine relative TXNIP mRNA level (normalized to β-actin) and the levels of the indicated proteins in each cell population. The experiments shown in B and E were repeated at least twice and representative experiments are shown. Values are reported as mean ± sd. Statistical significance was determined using t tests. ***P < 0.001.

To overcome this hurdle, we stably expressed the human TXNIP coding region in MondoA knockout (MondoA KO) MEFs under the control of a constitutive promoter (i.e. control cells) and examined effects of Ras activation by expressing Ras^G12V^ (i.e. Ras^G12V^-expressing cells). All subsequent experiments were conducted using these cells, unless otherwise specified. MondoA KO MEFs lack endogenous TXNIP expression, enabling investigation of TXNIP expression from the exogenous TXNIP allele (Fig.1D). In this experimental system, the level of ectopic TXNIP mRNA was higher in Ras^G12V^-expressing cells than in control cells (Fig.1E), yet TXNIP protein expression was dramatically repressed by Ras^G12V^ (Fig.1F). This experiment suggests that Ras activation induces TXNIP protein degradation or suppresses TXNIP synthesis.

### Ras^G12V^ inhibits TXNIP translation

We determined whether Ras^G12V^ increases the rate of TXNIP protein degradation using two methods. We first measured TXNIP protein degradation by blocking *de novo* protein synthesis with cycloheximide (CHX). We observed that the half-life of TXNIP protein was about 1 h in control cells and shorter (about 40 min) in Ras^G12V^-expressing cells, suggesting that Ras activation can increase TXNIP turnover (Fig.2A and B). Consistent with this finding, the proteasome inhibitor MG132 also increased TXNIP protein levels in Ras^G12V^-expressing cells, but the increase was not to the levels observed in control cells (Fig.2C). Collectively, these results suggest that Ras^G12V^ stimulates TXNIP degradation, but this increase in degradation rate does not fully account for the difference in TXNIP levels in control and Ras^G12V^-expressing cells.

**FIG 2.**
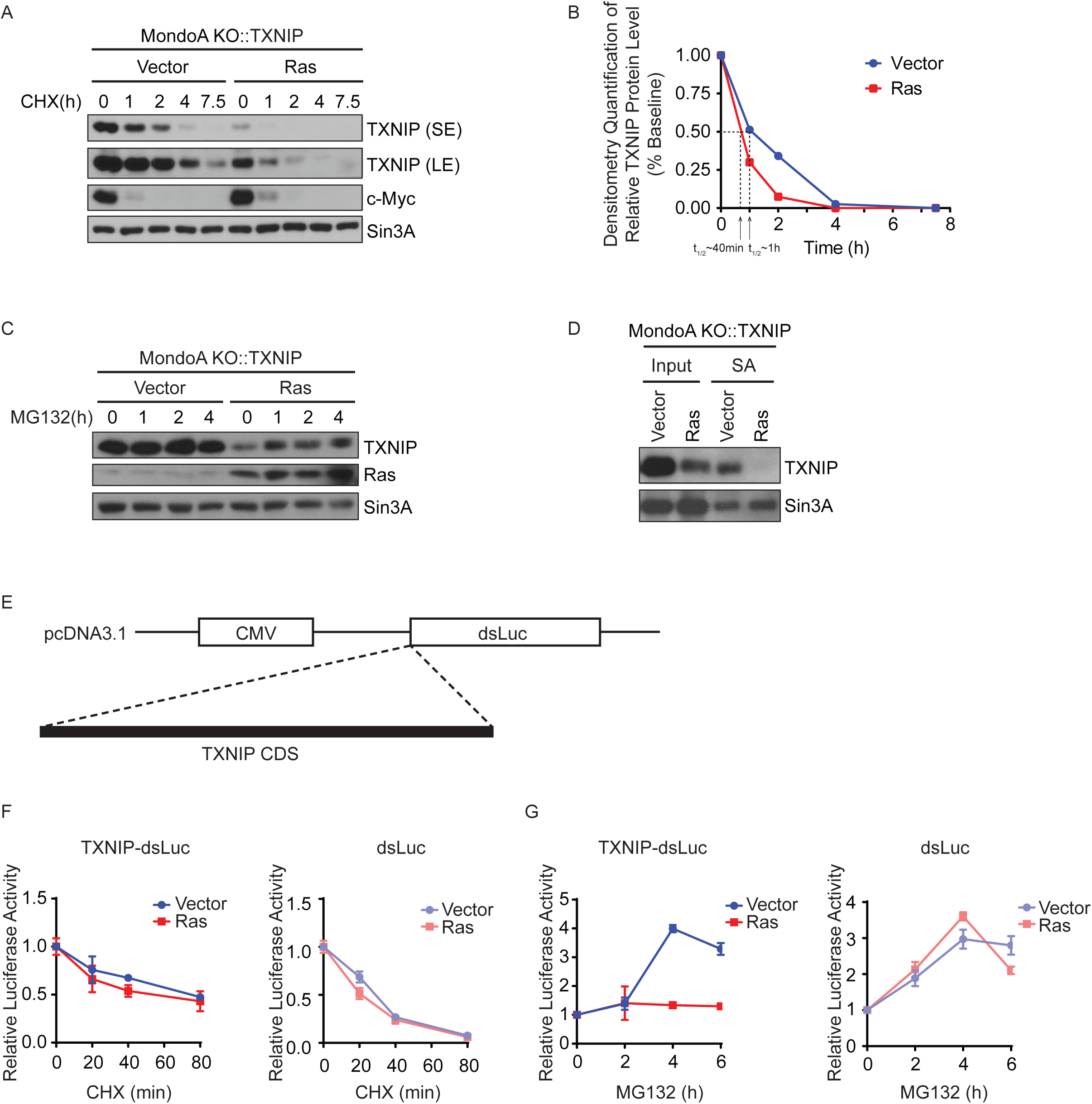
Ras^G12V^ inhibits TXNIP translation. **(A)** Western blotting was used to determine the levels of the indicated proteins in the listed cell lines following a treatment time course with 40 μg/ml cycloheximide (CHX). Short (SE) and long (LE) exposures allow better visualization of the TXNIP signal. c-Myc and Sin3A serve as controls for proteins that turnover rapidly or slowly. **(B)** TXNIP protein levels in (A) were quantified using densitometry. Signals from short exposure (SE) or long exposure (LE) were used for quantification for the control cells (Vector) or the Ras^G12V^-expressing cells (Ras), respectively. The half-life (t_1/2_) of TXNIP protein in each cell line was determined and indicated in the plot. **(C)** Western blotting was used to determine the levels of the indicated proteins in the listed cell lines following a treatment time course with 20 μM MG132. **(D)** Newly-synthesized proteins were enriched by labeling cells with azidohomoalanine (AHA), followed by biotinylation using Click chemistry and affinity purification using streptavidin beads (SA). Western blotting was used to determine the levels of steady-state (Input) and newly-synthesized (SA) TXNIP and Sin3A proteins in the indicated cell lines. **(E)** Diagram of the pcDNA3.1-TXNIP-dsLuc2CP construct in which the human TXNIP CDS was cloned upstream of and in frame with the destabilized luciferase ORF (dsLuc). **(F-G)** Luciferase reporters were transfected into MondoA KO::TXNIP::Vector (Vector) or Ras^G12V^ (Ras) MEFs. Relative luciferase activity, normalized to β-gal activity, of TXNIP-dsLuc or dsLuc was determined following treatment time courses with 40 μg/ml cycloheximide (CHX) (F) or 20 μM MG132 (G) in the indicated cell lines. The luciferase experiments were repeated at least twice and representative experiments are shown. Values are reported as mean ± sd.

A number of previous publications focused on TXNIP degradation (10, 29); therefore, we elected to investigate whether Ras^G12V^ affects the rate of TXNIP protein synthesis. Newly-synthesized proteins were identified by labeling cells with a methionine analog (azidohomoalanine, AHA), followed by their subsequent biotinylation using Click chemistry and enrichment on streptavidin beads. To eliminate potential effects from degradation, cells were also treated with MG132. As expected, Ras^G12V^-expressing cells had a lower level of steady-state TXNIP than control cells. Furthermore, Ras^G12V^-expressing cells expressed significantly less newly-synthesized TXNIP (Fig.2D). These data suggest that Ras^G12V^ suppresses TXNIP protein synthesis.

We next designed a luciferase reporter assay to confirm whether Ras^G12V^ suppresses TXNIP expression by blocking its synthesis. In this assay, the TXNP CDS was fused upstream of and in frame with the open reading frame encoding destabilized luciferase (dsLuc). To minimize transcriptional effects, dsLuc expression was driven from a constitutive promoter. Thus, the translation efficiency through the TXNIP CDS can be assessed by the luciferase activity (Fig.2E). In the presence of CHX, the activity of TXNIP-dsLuc and dsLuc decreased with a similar half-life in both control and Ras^G12V^-expressing cells, demonstrating that Ras^G12V^ does not affect the degradation of dsLuc or TXNIP-dsLuc (Fig.2F). By contrast, in the presence of MG132, the activity of TXNIP-dsLuc was significantly lower in Ras^G12V^-expressing cells compared to control cells, whereas Ras^G12V^ did not block the increase in dsLuc activity (Fig.2G). Together, these experiments suggest that Ras^G12V^ suppresses TXNIP expression primarily by decreasing TXNIP synthesis with an increase in TXNIP degradation being a secondary contributing factor. Furthermore, the TXNIP CDS is sufficient to confer the Ras^G12V^-dependent blockade of TXNIP synthesis.

### Ras^G12V^ inhibits translation elongation of TXNIP mRNA

Growth factor signaling promotes global protein synthesis (30, 31). Therefore, it is paradoxical that Ras^G12V^ suppresses TXNIP synthesis. To explore this contradiction further, we first determined whether Ras^G12V^ enhances global protein synthesis in MEFs using AHA labeling. As expected, global protein synthesis was increased after Ras activation (Fig.3A). Thus, even though Ras activation increases global translation, it appears to suppress translation of the TXNIP mRNA.

**FIG 3.**
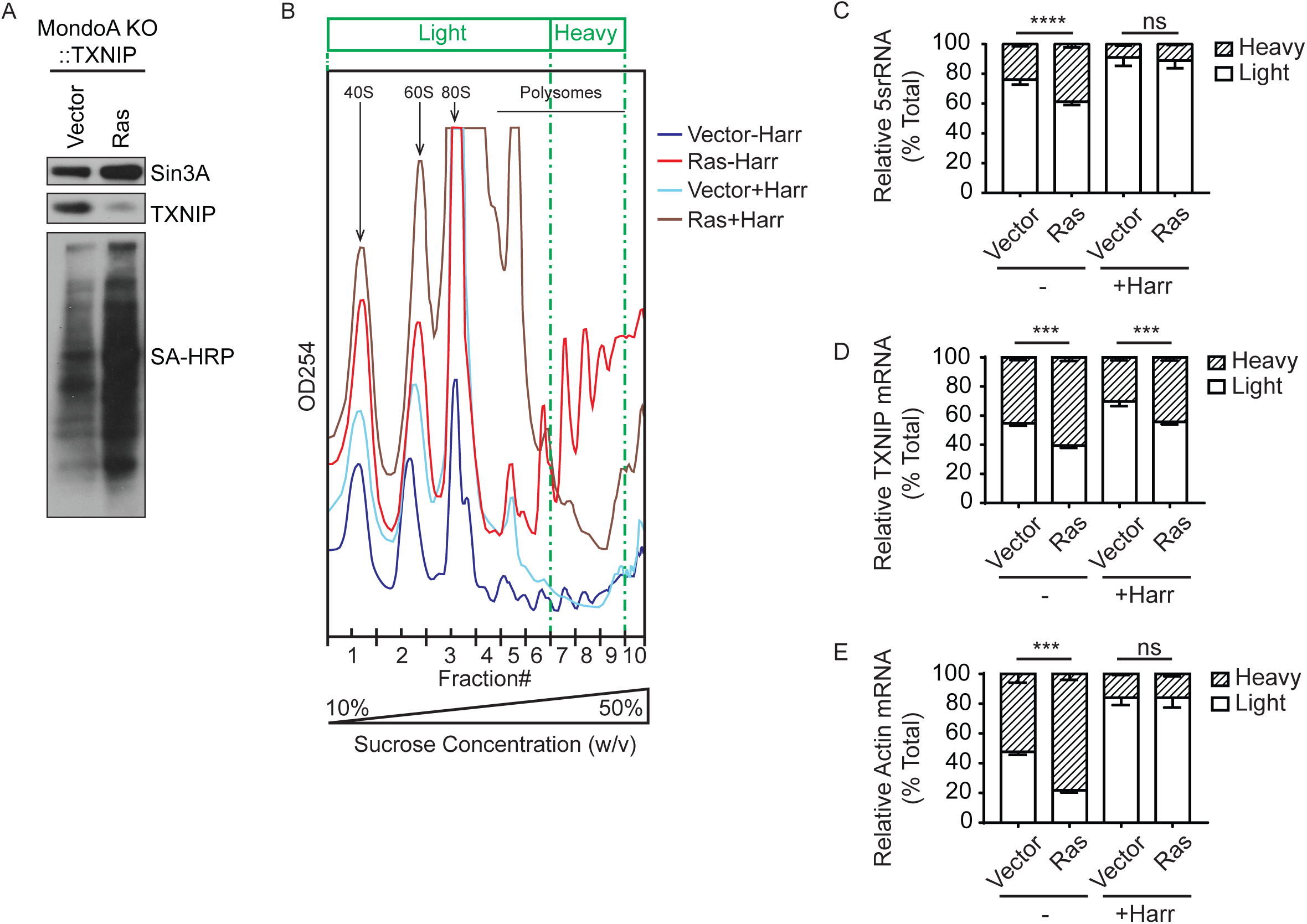
Ras^G12V^ inhibits translation elongation of TXNIP mRNA. **(A)** Cells were labeled with azidohomoalanine (AHA). Click chemistry was used to biotinylate the labeled proteins. Western blotting was used to determine the levels of TXNIP and Sin3A proteins and newly-synthesized proteins (SA-HRP) in the indicated cell lines. **(B)** Chromatograms showing the polysome profile of MondoA KO::TXNIP::Vector (Vector) or Ras^G12V^ (Ras) MEFs before (-Harr) and after (+Harr) harringtonine treatment. Fraction numbers are labeled on the x-axis; the post-qPCR pooling scheme is indicated by vertical dotted lines and labeled on the top of the graph (Light fraction: mRNAs containing <=3 ribosomes; Heavy fraction: mRNAs containing >=4 ribosomes). **(C-E)** RT-qPCR was used to determine the amount of 5srRNA (C), TXNIP mRNA (D), and Actin mRNA (E) in the heavy and light fractions from the polysome profile of the listed cell population (see 3B). The amount of RNAs in the heavy and light fractions is normalized to total RNA. Experiments were repeated twice and a representative experiment is shown. Values are reported as mean ± sd. Statistical significance was determined using one-way ANOVA. ***P < 0.001; ****P < 0.0001; ns P ≥ 0.05.

**FIG 4.**
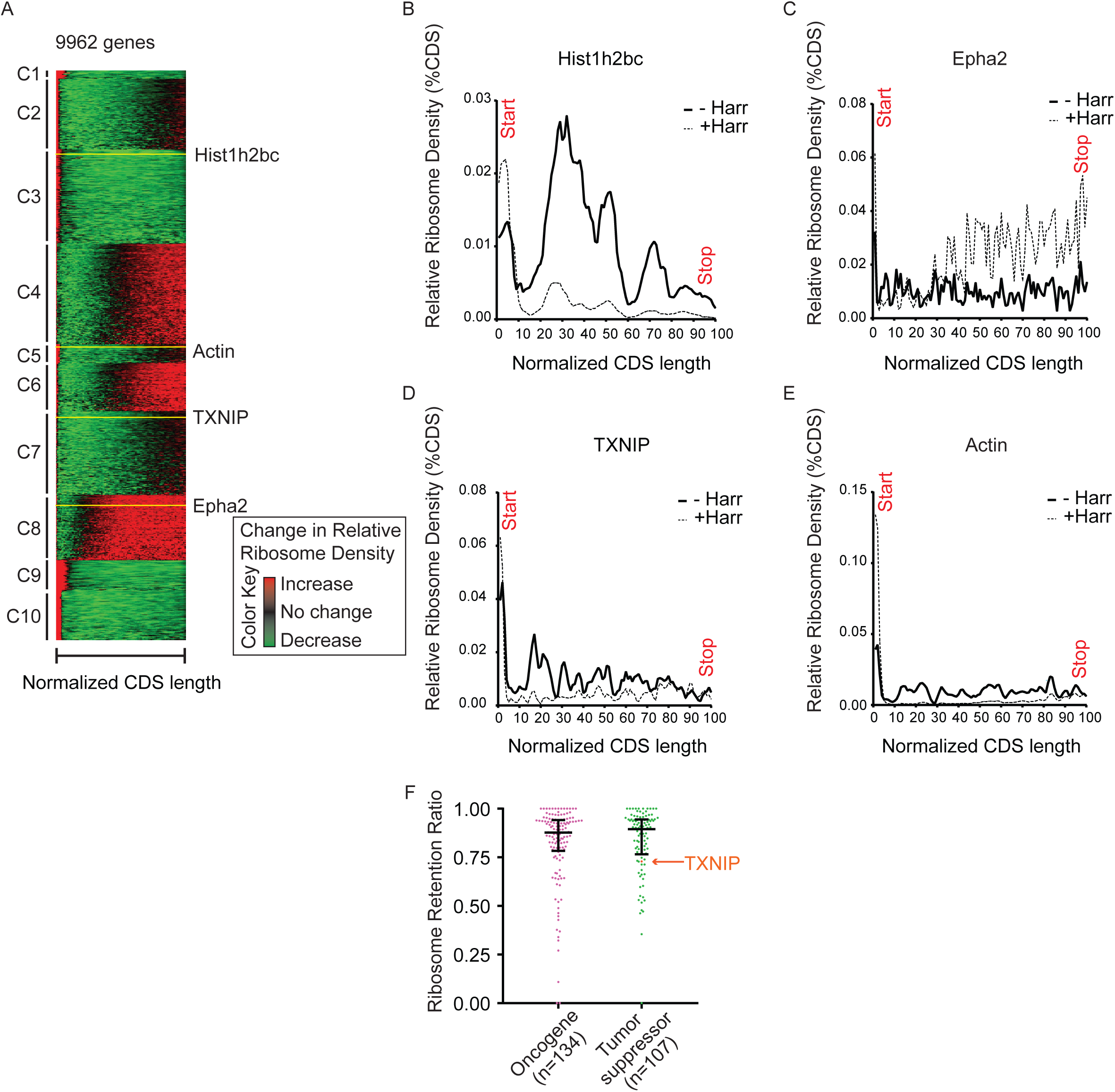
Translation elongation rate does not correlate with gene function. **(A)** Heatmap showing unsupervised clustering on the top 9962 translationally active gene transcripts based on their elongation profile, which was generated by calculating the change of relative ribosome density across the CDS of each transcript caused by harringtonine treatment. Red and green indicate increased and decreased relative ribosome density after harringtonine treatment, respectively. The Hist1h2bc, Epha2, TXNIP and Actin transcripts are indicated by yellow lines. **(B-E)** Ribosome profiling data showing the relative ribosome density across the CDS of Hist1h2bc, Epha2, TXNIP, and Actin transcripts in mECSs before (-Harr) and after (+Harr) harringtonine treatment. **(F)** Grouped scatter plot showing the ribosome retention ratio of 134 oncogenes and 107 tumor suppressors, with median and interquartile range indicated on the plot.

**FIG 5.**
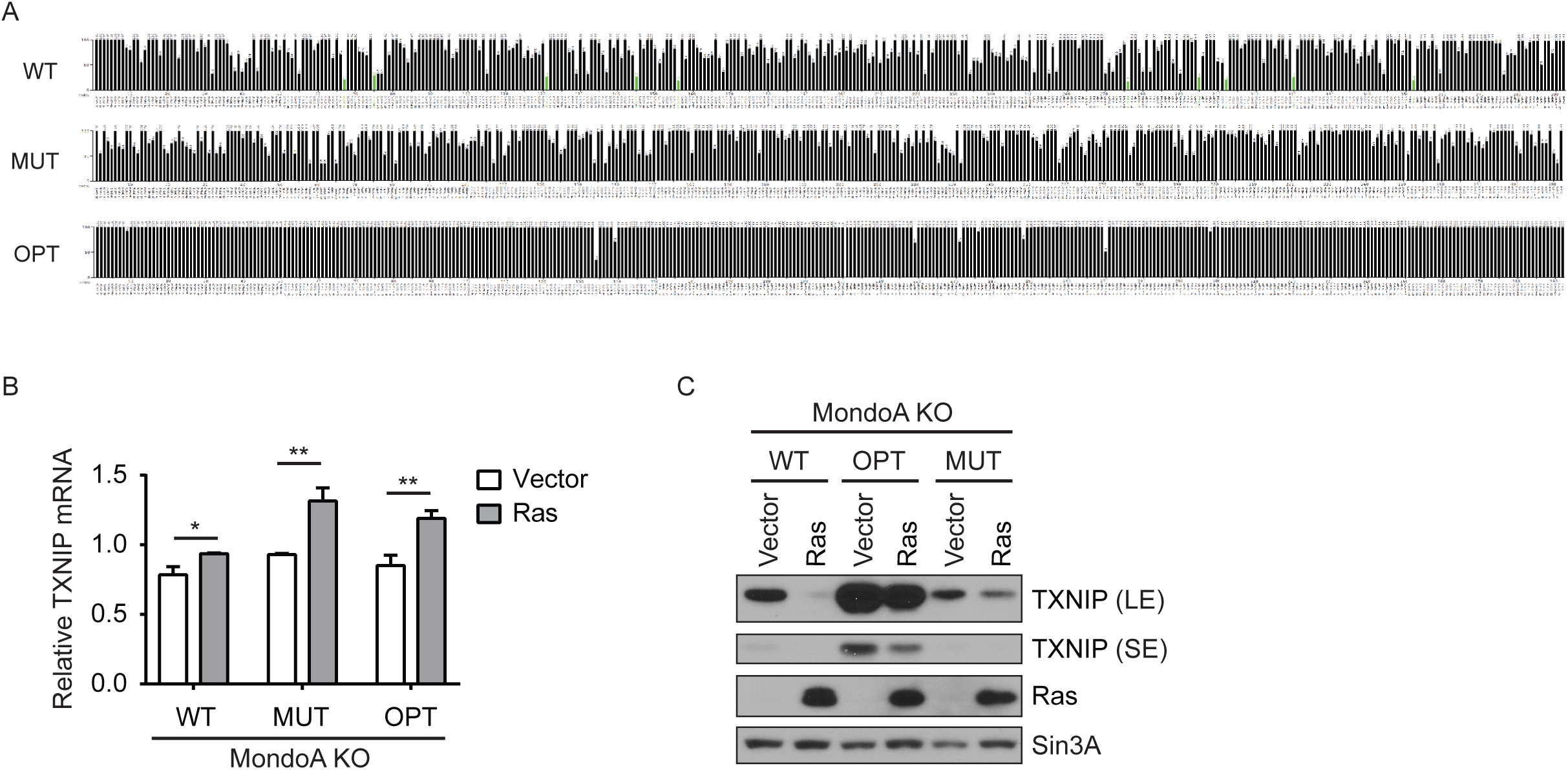
TXNIP mRNA primary sequence is not required for Ras^G12V^-dependent translational repression. **(A)** Diagram showing relative adaptiveness of each codon of TXNIP-WT, -MUT and -OPT. **(B-C)** MondoA KO MEFs were stably transduced with TXNIP variants (WT, MUT, or OPT), each of which was further stably transduced with pBabePuro (Vector) or pBabePuro-H-Ras^G12V^ (Ras) as indicated. RT-qPCR (B) and western blotting (C) were used to determine relative TXNIP mRNA level (normalized to β-actin) and the levels of the indicated proteins in each cell population. Long (LE) and short (SE) exposures allow better visualization of the TXNIP signal. Values in B are reported as mean ± sd. Statistical significance was determined using t test. *P < 0.05; **P < 0.01.

**FIG 6.**
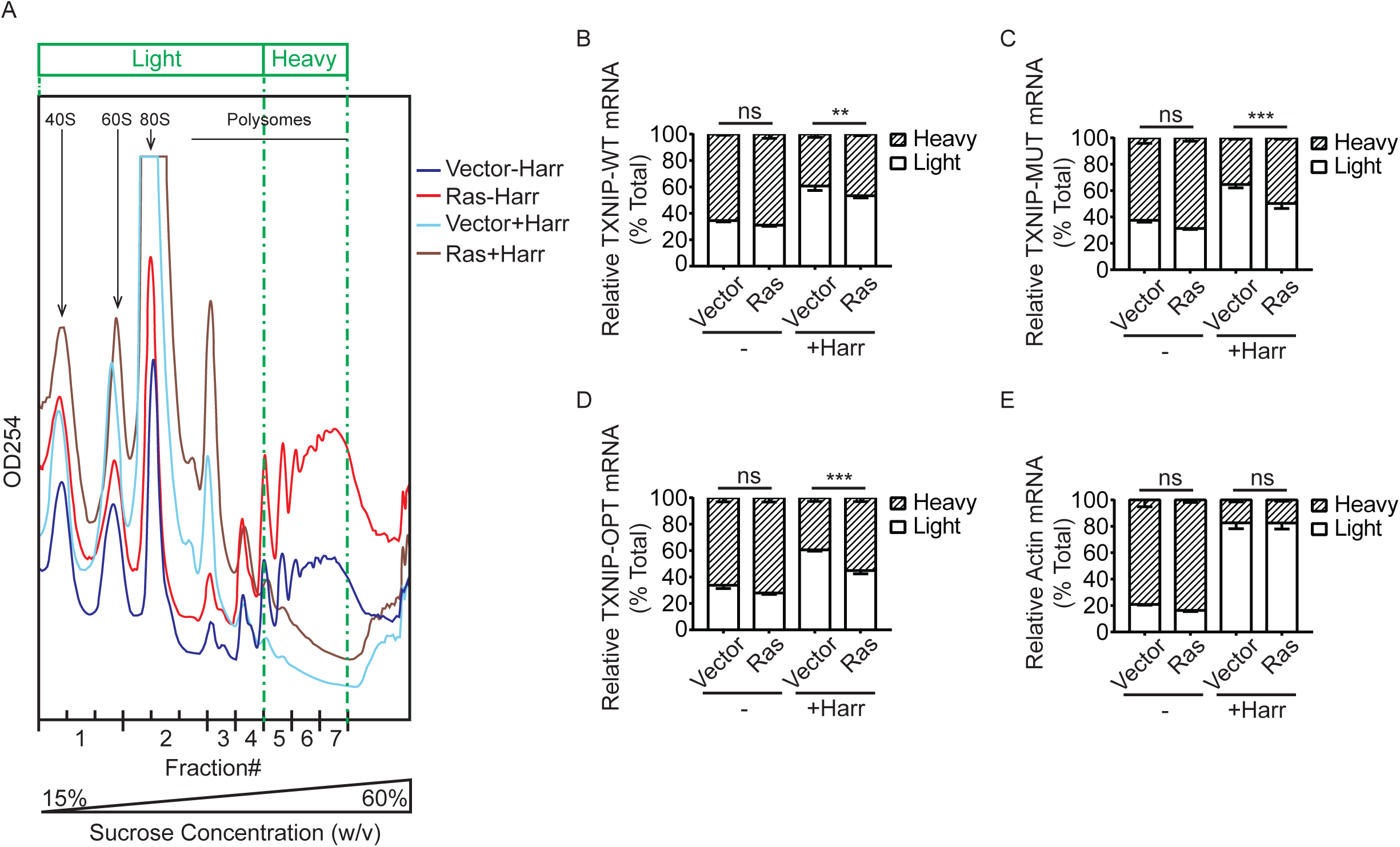
Ras^G12V^ inhibits TXNIP translation elongation independent of TXNIP’s mRNA primary sequence. **(A)** Chromatograms showing the polysome profile of MondoA KO::TXNIP variants::Vector (Vector) or Ras^G12V^ (Ras) MEFs before (-Harr) and after (+Harr) harringtonine treatment. Fraction numbers are labeled on the x-axis; the post-qPCR pooling scheme is indicated by vertical dotted lines and labeled on the top of the graph (Light fraction: mRNAs containing <=3 ribosomes; Heavy fraction: mRNAs containing >=4 ribosomes). **(B-E)** RT-qPCR was used to determine the amount of TXNIP-WT mRNA (B), TXNIP-MUT mRNA (C), TXNIP-OPT mRNA (D), and Actin mRNA (E) in the heavy and light fractions from the polysome profile of the listed cell population (see 6A). The amount of RNAs in the heavy and light fractions is normalized to total RNA. Experiments were repeated twice and a representative experiment is shown. Values are reported as mean ± sd. Statistical significance was determined using one-way ANOVA. **P < 0.01; ***P < 0.001; ns P ≥ 0.05.

**FIG 7.**
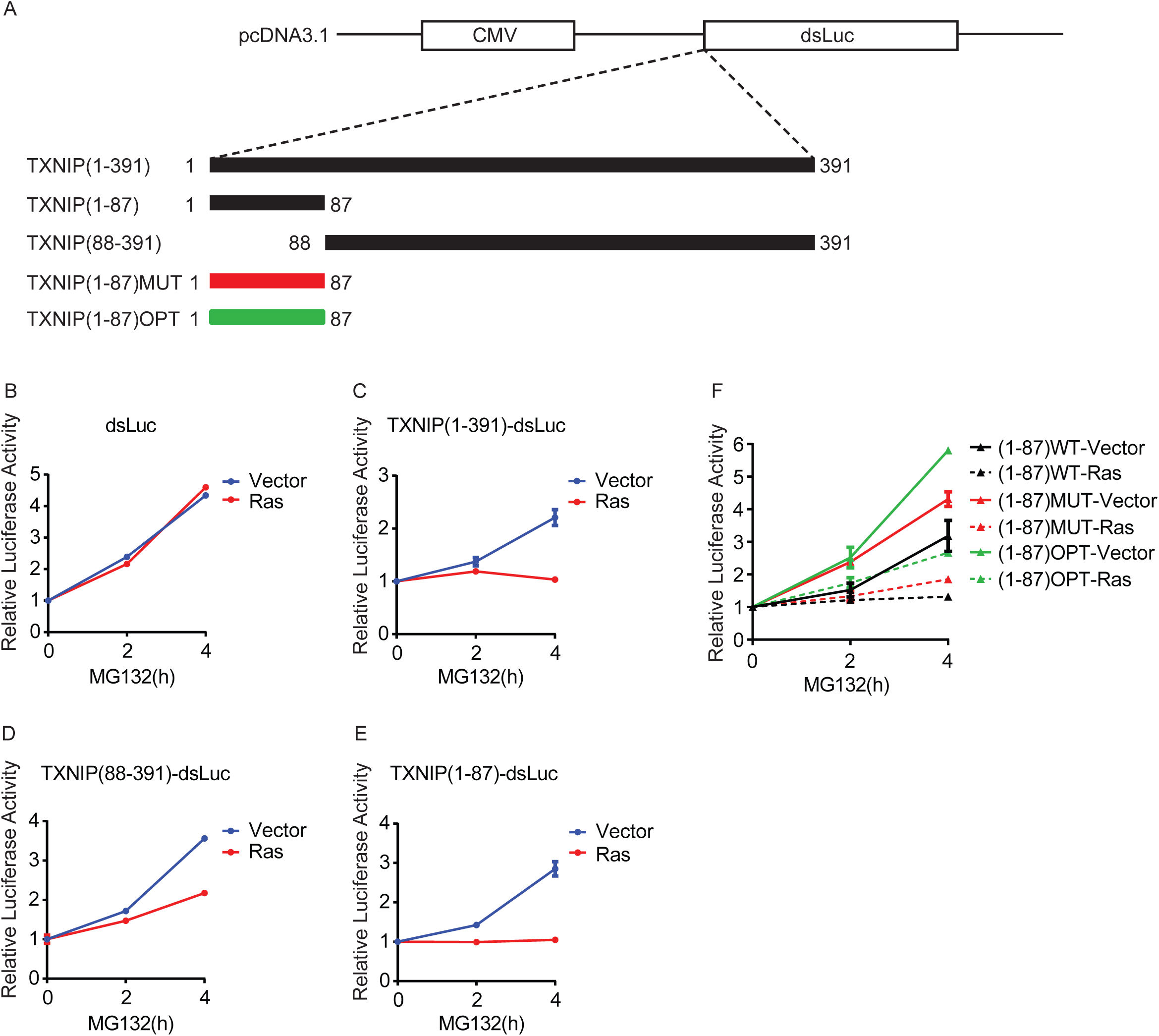
TXNIP N-terminal peptide sequence mediates the translational repression by Ras^G12V^. **(A)** Diagram showing luciferase reporter constructs, in which the human full-length TXNIP(1-391), TXNIP(1-87), TXNIP(88-391), TXNIP(1-87)MUT or TXNIP(1-87)OPT was cloned upstream of and in frame with the destabilized luciferase ORF (dsLuc). **(B-F)** Luciferase reporters were transfected into MondoA KO::TXNIP::Vector (Vector) or Ras^G12V^ (Ras) MEFs. Relative luciferase activity, normalized to β-gal activity, of each reporter was determined following a treatment time course with 20 μM MG132 in the indicated cell lines. The luciferase experiments were repeated at least twice and representative experiments are shown. Values are reported as mean ± sd.

Since the TXNIP CDS is sufficient for translational repression by Ras^G12V^ (Fig.2D and G), we hypothesized that Ras^G12V^ restricts TXNIP synthesis by blocking translation elongation of the TXNIP message. To test this hypothesis, we used polysome profiling to determine how Ras^G12V^ affects the distribution of ribosomes on the TXNIP mRNA and other messages. To gauge elongation rate, we used harringtonine, which is a translation initiation inhibitor that immobilizes initiating ribosomes at the start codon without affecting elongating ribosomes. Thus, the rate at which ribosomes are cleared from individual mRNAs in the presence of harringtonine can be used to estimate the elongation rate (32, 33). Cell lysates from control and Ras^G12V^-expressing cells were subject to velocity sedimentation in a sucrose gradient and the gradient was subsequently fractionated. RNA was purified from each fraction and the levels of gene transcripts of interest were determined by RT-qPCR. To simplify the analysis, qPCR values obtained for each fraction were combined into “light” vs “heavy” bins and presented as a percentage of the total RNA (Light fraction: mRNAs containing <=3 ribosomes; Heavy fraction: mRNAs containing >=4 ribosomes) (Fig.3B).

We observed that Ras^G12V^-expressing cells had significantly more polysome-associated mRNA present in the heavy fraction than control cells, confirming that Ras activation drives global translation. Moreover, the majority of mRNA shifted from the heavy fraction to the light fraction in the presence of harringtonine in both cell types, suggesting effective clearance of ribosomes from the mRNA transcripts (Fig.3B). We first examined 5srRNA, which reflects global ribosome association of mRNAs. Consistent with the chromatogram, 5srRNA was more enriched in the heavy fraction from Ras^G12V^-expressing cells than from control cells. After harringtonine treatment, 5srRNA shifted to the light fraction to the same extent for both cell populations (Fig.3C). TXNIP mRNA was more enriched in the heavy fraction from Ras^G12V^-expressing cells than from control cells. However, TXNIP mRNA was retained in the heavy fraction from Ras^G12V^-expressing cells after the harringtonine treatment, suggesting that active Ras slows the rate at which ribosomes transit the TXNIP message (Fig.3D). For comparison, we examined Actin mRNA, a housekeeping gene expected to be translated efficiently. Actin mRNA was more enriched in the heavy fraction from Ras^G12V^-expressing cells and shifted to the light fraction for both cell types upon harringtonine treatment (Fig.3E).

### Translation elongation rate does not correlate with gene function

We wondered whether Ras’s blockade of elongation is restricted to TXNIP or does it suppress translation more broadly. To investigate this question, we analyzed data from a ribosome profiling experiment designed to examine translation elongation in murine embryonic stem cells (33). We generated the elongation profile for each gene transcript by calculating the change of relative ribosome density across the CDS caused by harringtonine treatment. We next performed unsupervised clustering on the top 9962 translationally active gene transcripts based on their elongation profile. The 9962 genes were segregated into 10 clusters (C1-C10) with a wide dynamic range. For example, ribosomes on transcripts from C1, C3, C9 and C10 elongate extremely fast while ribosomes on transcripts from C4, C6 and C8 elongate extremely slowly (Fig.4A). An example of a “fast” gene is Hist1h2bc in C3, whereas an example of a “slow” gene is Epha2 in C8 (Fig.4B and C). Consistent with our polysome profiling data, ribosomes run off the TXNIP message, in C7, relatively slowly whereas they run off the Actin message, in C5, relatively quickly (Fig.4D and E). We performed pathway analysis for the genes in C7 and found that this cluster is not enriched for growth suppressors. Therefore, messages with elongation dynamics similar to TXNIP do not generally encode growth suppressors. Further, this finding suggests that Ras suppression of translation elongation might be restricted to TXNIP mRNA.

Even though growth suppressors did not cluster with TXNIP in C7, we wondered whether tumor suppressors and oncogenes have different elongation dynamics, allowing differential translational regulation by Ras or other oncogenic lesions. By comparing the ribosome retention ratio of 134 oncogenes and 107 tumor suppressors, we observed that the two groups have similar elongation dynamics (Fig.4F). Together, these results suggest that rapid elongation dynamics are not generally associated with messages encoding oncogenes, nor are slow elongation dynamics associated with messages encoding tumor suppressors.

### TXNIP mRNA primary sequence is not required for Ras^G12V^-dependent translational repression

We next investigated whether the sequence of the TXNIP message contributed to the Ras^G12V^-dependent blockade of translation elongation. First, we tested the contribution of the primary sequence of the TXNIP mRNA. We used gene synthesis to create a TXNIP mRNA with extensive silent mutations across the entire CDS. Furthermore, this artificial message, TXNIP-MUT, is comprised of codons with similar usage frequencies in the mouse genome to those of the TXNIP-WT mRNA (Fig.5A). In total, we altered 422 of 1176 bases of the TXNIP CDS without changing the TXNIP primary amino acid sequence. As expected, TXNIP-WT and TXNIP-MUT mRNA levels were comparable in control cells and were increased upon Ras activation (Fig.5B). Nevertheless, TXNIP-MUT was still repressed by Ras^G12V^, although to a lesser extent than TXNIP-WT (Fig.5C). This suggests that the translational repression of TXNIP by Ras^G12V^ is unlikely to be mediated by a feature(s) of the primary sequence of the TXNIP mRNA, such as mRNA secondary structure, miRNAs or sequence-specific RNA binding proteins.

The WT TXNIP mRNA is comprised of a significant number of sub-optimal codons and even some rare codons. Therefore, we speculated that WT TXNIP mRNA might be at a competitive disadvantage for translation machinery in cells where Ras^G12V^ drives global translation. We again used gene synthesis to generate an artificial TXNIP mRNA. In this case, we retained the coding capacity of the TXNIP mRNA, but replaced sub-optimal codons with codons that are used most frequently in the mouse genome. In total, we altered 240 of 1176 bases (Fig.5A). This mRNA, TXNIP-OPT was transcribed similarly to TXNIP-WT (Fig.5B). Yet, the level of TXNIP protein encoded by the TXNIP-OPT mRNA was much higher than that expressed from the TXNIP-WT mRNA, confirming that high frequency codons can increase the translation efficiency. However, TXNIP-OPT was still subject to translational suppression by Ras^G12V^ (Fig.5C). This finding suggests that Ras activation does not suppress translation of the TXNIP mRNA simply because it contains sub-optimal or even rare codons. Further, the high level of mutation in the TXNIP-OPT mRNA also supports the model that the primary mRNA sequence is not targeted by Ras^G12V^.

### Ras^G12V^ inhibits TXNIP translation elongation independent of TXNIP’s mRNA primary sequence

Next, we investigated whether the primary sequence of the TXNIP mRNA is involved in the elongation repression by Ras^G12V^. To test this, we examined the elongation rate of TXNIP-WT, - MUT and -OPT messages with and without Ras activation using polysome profiling and the ribosome run-off assay (Fig.6A). In the absence of harringtonine, the loading of ribosomes onto all three TXNIP messages was identical in control and Ras^G12V^-expressing cells. However, after harringtonine treatment, each TXNIP message was more highly enriched in the heavy fraction in Ras^G12V^-expressing than in control cells (Fig.6B-D). As a control, Actin exhibited equally efficient run-off in control and Ras^G12V^-expressing cells (Fig.6E). These results further support the model that Ras^G12V^ suppresses translation elongation of the TXNIP mRNA independent of its primary sequence or codon usage.

### TXNIP N-terminal peptide sequence mediates the translational repression by Ras^G12V^

Together, our results suggest that a feature of the TXNIP protein sequence is targeted by Ras. Therefore, we investigated which region of the TXNIP protein is necessary or sufficient for the translational repression by Ras^G12V^. To achieve this goal, we fused different regions of TXNIP to dsLuc (Fig.7A). As above, Ras^G12V^ did not affect the accumulation of dsLuc activity but completely suppressed the accumulation of TXNIP-dsLuc activity after MG132 treatment (Fig.7B and C). However, TXNIP(88-391)-dsLuc activity was partially sensitive to Ras^G12V^ suppression, suggesting a critical role for the first 87 amino acids of TXNIP in Ras^G12V^-dependent translational repression (Fig.7D). Consistent with this, Ras^G12V^ completely suppressed the activity of TXNIP(1-87)-dsLuc (Fig.7E). This finding suggests that the first 87 amino acids of TXNIP are sufficient for the Ras^G12V^-dependent repression. To test the potential contribution of mRNA primary sequence and/or codon frequency in this context, we fused artificial TXNIP mRNAs that comprise the N-terminal 87 amino acids encoded by the MUT or OPT TXNIP sequences to dsLuc (Fig.5A and 7A). Consistent with our previous finding showing that Ras^G12V^ targets the TXNIP protein sequence, the activity of TXNIP(1-87)MUT-dsLuc and TXNIP(1-87)OPT-dsLuc were both repressed by Ras^G12V^ (Fig.7F).

## DISCUSSION

Our lab previously showed that acute growth factor signaling leads to a dramatic and rapid suppression of TXNIP expression. In that study, the reduction in TXNIP protein preceded the reduction in TXNIP mRNA, suggesting that growth factor signaling impacts TXNIP protein synthesis and/or degradation (27). In this study, we show that Ras activation suppresses TXNIP mRNA and protein expression. By bypassing transcriptional regulation, we show that Ras actively represses translation elongation of TXNIP mRNA. Translating ribosomes continue to transit the TXNIP message (Fig.3 and Fig.6), albeit at a slower rate in the presence of Ras activation; therefore, we suggest that Ras^G12V^ slows the rate of elongation rather than causing ribosome stalling. Ras activation also stimulates TXINIP turnover, but this mechanism only accounts for fraction (~25%) of the effect of Ras activation on TXNIP levels: the repression of TXNIP synthesis accounts for the majority of the remaining Ras-dependent decrease in TXNIP (Fig.2). Ras activation represses transcription of the TXNIP gene and controls TXNIP levels post-transcriptionally by inducing degradation of the TXNIP protein and blocking translation elongation of the TXNIP message. We speculate that this multi-faceted downregulation of TXNIP expression by activated Ras helps ensure that adequate glucose is available to support Ras-driven anabolic biosynthetic pathways.

Growth factor signaling is known to promote global protein synthesis, through upregulation of translation initiation and elongation (30, 31); therefore, it is striking that Ras activation blocks translation elongation on the TXNIP message. Like growth factor signaling, we show that Ras^G12V^ upregulates global protein synthesis in MEFs (Fig.3A). However, Ras activation blocks TXNIP synthesis. Our analysis of published polysome profiling data suggests that messages with similar elongation dynamics to TXNIP do not generally encode growth/tumor suppressors (Fig.4). This finding suggests that Ras^G12V^-mediated translational inhibition might be restricted to TXNIP or a smaller set of messages.

We next investigated the mechanisms by which Ras^G12V^ specifically downregulates the translation of TXNIP mRNA. The artificial TXNIP mRNAs, TXNIP-MUT and TXNIP-OPT, are both significantly different from TXNIP-WT in primary sequence. TXNIP-MUT is designed to alter the coding sequence without changing the overall codon usage frequency. Ras activation suppresses TXNIP expression when it is encoded by TXNIP-MUT. Therefore, we propose that Ras regulates TXNIP synthesis by a mechanism independent of mRNA primary sequence. TXNIP-OPT is designed with optimal codon usage, i.e. with increased usage of high frequency codons, which positively impacts the elongation rate (34-37). TXNIP-OPT is still subject to Ras^G12V^-dependent translational repression, suggesting that suboptimal codon usage in the TXNIP message is not responsible for the translational repression by Ras^G12V^. Supporting our model that the primary sequence of the TXNIP mRNA is not targeted by Ras^G12V^, the rate at which ribosomes elongate on TXNIP-MUT and TXNIP-OPT is much slower in the presence of Ras^G12V^. Together, these experiments suggest that Ras does not target the primary sequence of the TXNIP mRNA to repress TXNIP synthesis.

In contrast to the lack of involvement of the sequence of TXNIP mRNA, we discovered that the N-terminus of the TXNIP protein is necessary and sufficient for translational repression by Ras^G12V^. We note that the effect of deleting the N-terminus of TXNIP is only partial (Fig.7D), suggesting that other regions of TXNIP also contribute to the Ras-dependent blockade of TXNIP synthesis. Using TXNIP truncation variants TXNIP(1-87)MUT and TXNIP(1-87)OPT, we show that the peptide sequence rather than the mRNA sequence coding the first 87 amino acids is required for translational repression. These data suggest that activated Ras targets the N-terminus of TXNIP as it exits the ribosome, resulting in a decreased rate of translation elongation (Fig.8). The dependence on the N-terminus of TXNIP for Ras-dependent repression is consistent with our contention that the blockade of translation elongation by Ras may be restricted to TXNIP or to a small group of proteins that show homology to the TXNIP N-terminus.

**FIG 8.**
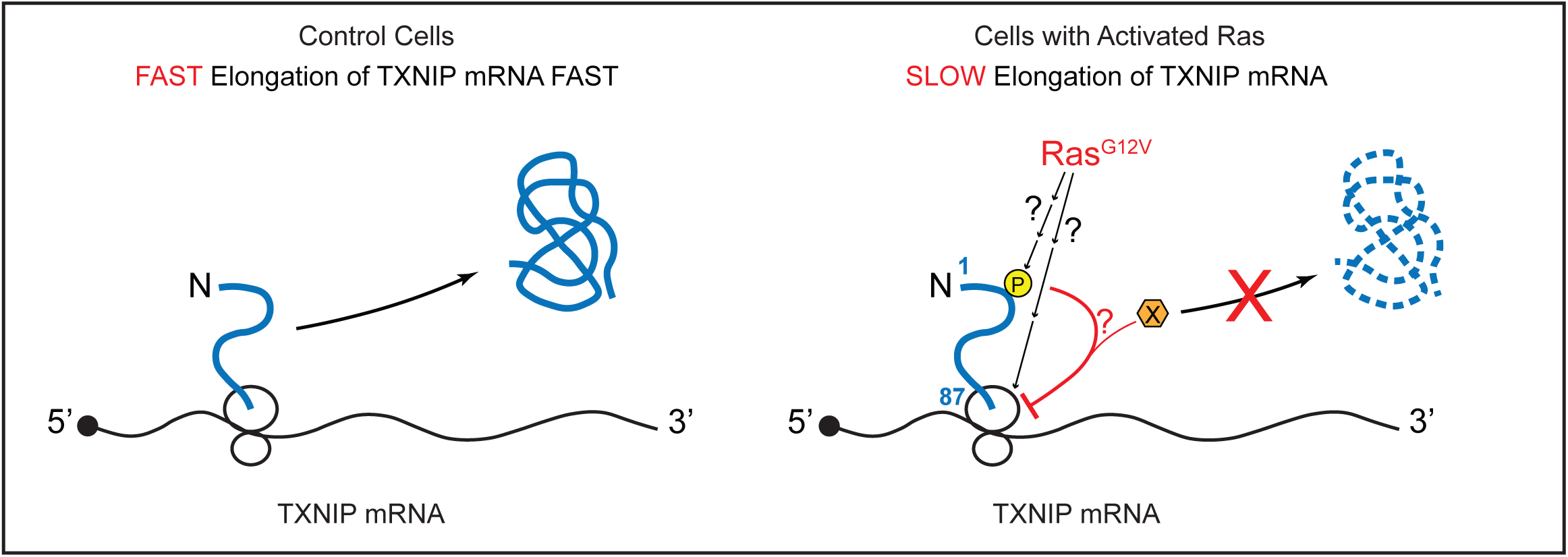
Ras^G12V^ suppresses TXNIP translation elongation. In unstimulated cells, a significant amount of TXNIP protein is synthesized from fast translation elongation of the TXNIP mRNA. In Ras-activated cells, TXNIP synthesis is suppressed due to the reduced rate at which ribosomes translate the TXNIP mRNA, via mechanism(s) that involve(s) the nascent peptide of TXNIP N-terminus. The first 87 amino acids of TXNIP protein or the ribosome itself (ribosomal proteins or rRNAs) might be targeted by Ras-dependent post-translational modifications, which impede peptide release, leading to slow elongation.

Accumulating evidence suggests that the nascent peptide chain regulates translation elongation by peptide sequence-specific interaction with the ribosome exit tunnel, which is accompanied by stalled ribosomal complexes (38-45). We observed two prominent peaks along TXNIP CDS corresponding to amino acids 66 and 81 of the TXNIP protein (Fig.4D), which is an indication of high ribosome density and potentially slow ribosome translocation. We propose that the slow elongation dynamics in this region of the TXNIP mRNA provides an opportunity for co-translational regulation by Ras^G12V^. We examined the ribosome distribution on ARRDC4 transcript, which is a TXNIP paralog that is also a potent negative regulator of glucose uptake (46). Interestingly, ARRDC4 transcript also shows high ribosome density at the position corresponding to amino acid 85 of ARRDC4 protein (data not shown), suggesting that its translation may also be regulated by Ras.

Ras^G12V^ targets the N-terminus of TXNIP, but we do not yet understand how that impacts translation elongation. Interaction between the nascent peptide chain and the ribosome exit tunnel affects the elongation dynamics (38, 39). In addition, co-translational modification of the protein being translated can regulate protein synthesis (47, 48). Given these findings, it is possible that Ras^G12V^ modifies the nascent peptide chain of TXNIP N-terminus, likely indirectly, to impede the peptide release and the elongation dynamics (Fig.8). Consistent with this idea, TXNIP harbors a number of conserved sites for potential post-translational modification within its first 87 amino acids. A second possibility comes from the observation that modifications of ribosomal proteins and rRNAs regulate translation in a growth phase-dependent manner and in response to environmental signals (49, 50). Therefore, it is also possible that Ras^G12V^ targets the ribosome itself, through affecting its interaction with the nascent peptide chain of TXNIP N-terminus, to achieve specific blockade of TXNIP elongation (Fig.8). Additional studies will be necessary to test these models and others. Our preliminary studies suggest that canonical Ras effector pathways do not block TXNIP translation (data not shown). Thus, Ras may drive metabolic reprogramming toward aerobic glycolysis by regulating translation elongation of the TXNIP mRNA, and potentially other messages, by a novel mechanism.

## MATERIALS AND METHODS

### Cell culture and conditions

All cells were maintained at 37°C in 5% CO_2_ in DMEM (Gibco11995-065) supplemented with 10% fetal bovine serum (FBS) (Gibco26140-079) and 1% penicillin/streptomycin (Gibco15140-122). Wildtype and MondoA knockout (MondoA KO) mouse embryonic fibroblasts (MEFs) were generated in the lab as described previously (23). MEFs were immortalized with SV40 large T antigen.

### Plasmids and virus infection

Retroviral constructs pBabePuro and pBabePuro-H-Ras^G12V^ were gifts from S. Lessnick. Human TXNIP variants (WT, MUT, and OPT) were cloned into pWzlBlast retroviral vector. TXNIP-MUT and TXNIP-OPT were generated by gBlocks gene fragments (Integrated DNA Technologies) and Gibson cloning (NEB E2621L). Sequences of TXNIP-MUT and TXNIP-OPT are available upon request. Retroviruses of each construct were packaged in HEK293EBNA cells with VSVG and gagpol plasmids. Stable cell lines were generated by retroviral infection followed by antibiotic selection (2 μg/ml puromycin for pBabePuro constructs; 3.5 μg/ml blasticidin for pWzlBlast constructs).

### Relative codon adaptiveness

Relative codon adaptiveness for TXNIP-WT, -MUT, and -OPT was calculated against mouse codon usage table and plotted using the web-based graphical codon usage analyzer (http://gcua.schoedl.de/sequential_v2.html).

### Chemicals and reagents

For proteasome inhibition, 20 μM MG132 (Calbiochem 474790) was used for the indicated times. For protein synthesis inhibition, 40 μg/ml cycloheximide (Sigma C7698) was used for the indicated times. For ribosome run-off assay, cells were treated with 2 μg/ml harringtonine for 10 min (Cayman Chemical 15361).

### Western blotting

Whole cell lysate was prepared in ice-cold lysis buffer (400 mM NaCl, 20 mM Hepes [pH7.6], 0.1% NP-40, 25% glycerol, 1 mM EDTA, 1 mM EGTA, 1 mM DTT) containing protease inhibitors (1 mM PMSF, 2.5 μg/ml aprotinin, 1 μg/ml leupeptin, 1 μg/ml pepstatin) and phosphatase inhibitors (phosphatase inhibitor cocktail 1 and 2, Sigma P2850 and P5726). Protein concentration was determined by Bio-Rad protein assay (Bio-Rad 500-0006). The same protein amount for individual samples were resolved on SDS-PAGE and transferred to PVDF membranes, and subsequently blocked in 5% non-fat milk in TBST (1x Tris-buffered saline with 0.1% Tween-20) followed by probing with primary antibodies at 1:2,000 dilution (MLXIP/MondoA, Proteintech 13614-1-AP; TXNIP, Abcam ab188865; H-Ras(c-20), Santa Cruz sc-520; c-Myc(Y69), Abcam ab32072) overnight at 4°C. HRP-conjugated mouse IgG (1:5,000) (GE Healthcare NA931V) or rabbit IgG (1:15,000) (GE Healthcare NA934V) and Western Lightning Plus-ECL (PerkinElmer NEL104001EA) were used for signal detection. Experiments were repeated at least twice and representative experiments are shown.

### Click assay

Cells were cultured with complete media lacking methionine (1x DMEM without L-Gln and L-Met, MP Biomedicals 1642254, supplemented with 4 mM Glutamine and 2% FBS) for 1 h prior to azidohomoalanine (AHA) labeling. MG132 (20 μM) was added to the -Met media 30 min before AHA labeling. AHA (Invitrogen C10102) was then added to the -Met media at a final concentration of 50 μM and cells were labeled for 3 h. Whole cell lysate was prepared in ice-cold RIPA buffer (50 mM Tris-HCl [pH7.4], 150 mM NaCl, 1% NP-40, 0.1% SDS, 0.25% Na deoxycholate) containing protease inhibitors (1 mM PMSF, 2.5 μg/ml aprotinin, 1 μg/ml leupeptin, 1 μg/ml pepstatin) and phosphatase inhibitors (50 mM NaF, 10 mM β-glycerophosphate, 1.5 mM Na_3_VO_4_). Protein concentration was determined by detergent compatible protein assay (Bio-Rad 500-0112). 100 μg of lysate was subject to Click reaction using biotin alkyne (Invitrogen B10185) and Click-iT protein reaction buffer kit (Invitrogen C10276) overnight at 4°C following manufacturer’s protocol.

For measurement of global protein synthesis, same volume of reacted lysate was analyzed by western blotting with HRP-conjugated streptavidin (1:2,000) (GE Healthcare RPN1231V). For measurement of TXNIP protein synthesis rate, the reacted lysate was chloroform/MeOH precipitated and resuspended in resolubilization buffer (60 mM Tris [pH6.8], 1% SDS, 10% glycerol). Protein concentration was determined using detergent compatible protein assay. 7.5 ug of the resolubilized lysates were subject to streptavidin (SA) affinity purification in RIPA buffer at 4°C overnight. Same amount of lysate (equivalent amount of SA-affinity purified samples as input) was subject to western blotting. Experiments were repeated at least twice and representative experiments are shown.

### Luciferase reporter assay

pcDNA3.1dsLuc2CP (dsLuc) was purchased from Addgene (#68054). TXNIP(1-391), TXNIP(1-87)-WT, TXNIP(1-87)-MUT, TXNIP(1-87)-OPT and TXNIP(88-391) were PCR amplified from respective pWzlBlast-TXNIP (WT, MUT, or OPT) constructs (as described above) with specific primers with 25 bp overhangs and subsequently cloned upstream of and in frame with dsLuc in the pcDNA3.1dsLuc2CP plasmid by Gibson cloning. Transfections were performed 24 h after cell plating when the cell monolayers were 60-70% confluent. Cells were treated with 40 μg/ml cycloheximide (CHX) or 20 μM MG132 for the indicated times 22-24 h post transfection and harvested in reporter lysis buffer (Promega E397A). Luciferase reporter assays were performed as previously described (51). Luciferase activity was normalized to β-gal activity and presented as a relative value to 0 h for both MG132 and CHX experiments. Experiments were repeated at least twice and representative experiments are shown. Values were reported as mean ± sd of three technical replicates.

### Reverse transcription quantitative PCR (RT-qPCR)

Total RNA was extracted using Quick-RNA miniprep kit (Zymo Research). cDNA was generated from 100-300 ng RNA using GoScript Reverse transcription kit (Promega). qPCR was performed using CFX Connect Real-time system and CFX manager software (Bio-Rad). Relative mRNA expression levels were determined from standard curves for each primer set, and were normalized to β-actin expression. Experiments were repeated at least twice and representative experiments are shown. Values were reported as mean ± sd of three technical replicates. Statistical significance was determined using t test. RT primer sequences are available upon request.

### Polysome profiling

10%-50% or 15%-60% (w/v) sucrose gradients in gradient buffer (15 mM Tris [pH7.4], 15 mM MgCl_2_, 100 mM KCl, 1 mM DTT, 100 μg/ml CHX) were prepared in 5 ml ultracentrifuge tubes (Beckman 326819) the night before experiment and stored at 4°C. Cells were washed and incubated with ice-cold PBS containing 100 μg/ml CHX for 2 min. Cells were scraped into Eppendorf tubes and centrifuged at 2,000xg for 4 min at 4°C. The cell pellet was collected and lysed in polysome lysis buffer (15 mM Tris [pH7.4], 15 mM MgCl_2_, 100 mM KCl, 1% Triton X-100, 1 mM DTT, 100 μg/ml CHX, 500 U/ml SuperaseIn, Invitrogen AM2696) containing protease inhibitors (1 mM PMSF, 2.5 μg/ml aprotinin, 1 μg/ml leupeptin, 1 μg/ml pepstatin) and phosphatase inhibitors (phosphatase inhibitor cocktail 1 and 2, Sigma P2850 and P5726). The cell lysate was loaded onto the top of a sucrose gradient and centrifuged at 47,000rpm for 1 h at 4°C in a SW55 Ti rotor (Beckman 342194) and centrifugation was stopped with no brake. The centrifuged sample was fractionated with a syringe pump coupled with UA-6 detector that continuously monitors OD254 values (Teledyne ISCO). 250 ul fractions were collected from the beginning of the 40S peak towards the end of the gradient. After fractionation, fractions corresponding to 40S, 60S and 80S peaks were combined and fraction numbers were assigned (refer to Fig.3B and Fig.6A for fraction numbers on the x-axis). The same volume of each fraction was taken and supplemented with 0.1 ng of luciferase mRNA (Promega), which served as spike-in control. RNA was extracted using TRIzol LS reagent (Invitrogen) and Direct-zol RNA miniprep kit (Zymo Research). cDNA was synthesized from the same volume of RNA using GoScript Reverse transcription kit (Promega). Levels for genes of interest were determined by RT-qPCR. Data were normalized to the luciferase RNA level to account for differences in RNA extraction and cDNA synthesis. For data presentation, qPCR values obtained from each fraction were combined into the light fraction (mRNAs containing <=3 ribosomes) and the heavy fraction (mRNAs containing >=4 ribosomes) (Fig.3B and Fig.6A). The values for the light/heavy fraction were presented as % total. Experiments were repeated twice and representative experiments are shown. Values were reported as mean ± sd of three technical replicates. Statistical significance was determined using one-way ANOVA.

For polysome profiling of cells expressing TXNIP-WT, MUT or OPT (Fig.6), each cell line (MondoA KO::TXNIP-WT::Vector; -MUT::Vector; -OPT::Vector; -WT::Ras^G12V^; -MUT:: Ras^G12V^; -OPT:: Ras^G12V^) for each treatment group (-Harr or +Harr) was plated and treated individually. During harvest, control cells with different TXNIP variants (WT, MUT or OPT) from the same treatment group were pooled (Vector-Harr or Vector+Harr); Similarly, Ras^G12V^-expressing cells with different TXNIP variants (WT, MUT or OPT) from the same treatment group were pooled (Ras-Harr or Ras+Harr). The sample pooling method was used to ensure identical experiment conditions. The resulting four samples were subject to polysome profiling and subsequent analysis to determine the RNA levels of genes of interest.

### Informatics

Gene expression values (mRNA expression z-Scores) of TXNIP and H-Ras-dependent genes (52) were obtained for the breast cancer (METABRIC) (53), lung adenocarcinoma (TCGA) (54) and pancreatic adenocarcinoma (TCGA, Provisional) datasets available on cBioPortal (55, 56) (http://www.cbioportal.org/). For the H-Ras gene signature, the z-Scores of the H-Ras signature genes were subject to principle component analysis (PCA) using the prcomp function in R (https://www.r-project.org/). Values of principle component 2 were used as H-Ras gene signature for each tumor sample.

Ribosome profiling data were obtained from the Gene Expression Omnibus (GEO) (Accession number GSE30839) (33). Sequencing data were preprocessed and aligned on the Galaxy server with the published method (57). To directly compare ribosome occupancy of mRNA transcripts across the entire genome, gene transcript length was normalized to 100 bins and relative ribosome density (% total reads from 100 bins) for each bin was calculated. Data were presented as relative ribosome density (y-axis) against normalized CDS length (x-axis). The elongation profile of each gene transcript was generated by calculating the difference of relative ribosome density caused by harringtonine treatment for each bin across the CDS. Unsupervised clustering was performed for the top 9962 translationally active gene transcripts based on their elongation profile using the Cluster3.0 program. The clustering result was visualized in heatmap using the TreeView program, in which red indicates an increase in relative ribosome density upon harringtonine treatment; whereas green indicates a decrease in relative ribosome density upon harringtonine treatment. The ribosome retention ratio (RRR) was calculated as the percentage of the cumulative relative ribosome density of the last 90 bins for each transcript in harringtonine-treated samples. An RRR of value closer to 0 indicates faster elongation rate; whereas an RRR of value closer to 1 indicates slower elongation rate. The lists of oncogenes and tumor suppressors were obtained from the UniProt database. The ribosome retention ratio of each gene was plotted as a grouped scatter plot, with median and interquartile range indicated on the graph.

## ACKNOWLEDGEMENTS

We thank the Weyrich lab for providing equipment and help for the polysome profiling experiments, the members of the Ayer lab for helpful discussion and insights, and Dr. Betty Leibold for comments on the manuscript. This work was supported by grants from the NIH, R01 GM055668-14A1 and the Department of Defense, BC133708, to D. E. A. Research reported in this publication utilized the High-Throughput Genomics and Bioinformatics Core at Huntsman Cancer Institute at the University of Utah and was supported by the National Cancer Institute of the National Institutes of Health under Award Number P30CA042014. The content is solely the responsibility of the authors and does not necessarily represent the official views of the NIH. The research reported in this publication was supported by Huntsman Cancer Foundation.

## REFERENCES

1. Bos JL. 1989. ras oncogenes in human cancer: a review. Cancer Res 49:4682–9.

2. Rajalingam K, Schreck R, Rapp UR, Albert S. 2007. Ras oncogenes and their downstream targets. Biochim Biophys Acta 1773:1177–95.

3. Hu Y, Lu W, Chen G, Wang P, Chen Z, Zhou Y, Ogasawara M, Trachootham D, Feng L, Pelicano H, Chiao PJ, Keating MJ, Garcia-Manero G, Huang P. 2012. K-ras(G12V) transformation leads to mitochondrial dysfunction and a metabolic switch from oxidative phosphorylation to glycolysis. Cell Res 22:399–412.

4. Gaglio D, Metallo CM, Gameiro PA, Hiller K, Danna LS, Balestrieri C, Alberghina L, Stephanopoulos G, Chiaradonna F. 2011. Oncogenic K-Ras decouples glucose and glutamine metabolism to support cancer cell growth. Mol Syst Biol 7:523.

5. Chiaradonna F, Sacco E, Manzoni R, Giorgio M, Vanoni M, Alberghina L. 2006. Ras-dependent carbon metabolism and transformation in mouse fibroblasts. Oncogene 25:5391–404.

6. Hu CJ, Wang LY, Chodosh LA, Keith B, Simon MC. 2003. Differential roles of hypoxia-inducible factor 1alpha (HIF-1alpha) and HIF-2alpha in hypoxic gene regulation. Mol Cell Biol 23:9361–74.

7. Kim JW, Zeller KI, Wang Y, Jegga AG, Aronow BJ, O’Donnell KA, Dang CV. 2004. Evaluation of myc E-box phylogenetic footprints in glycolytic genes by chromatin immunoprecipitation assays. Mol Cell Biol 24:5923–36.

8. Osthus RC, Shim H, Kim S, Li Q, Reddy R, Mukherjee M, Xu Y, Wonsey D, Lee LA, Dang CV. 2000. Deregulation of glucose transporter 1 and glycolytic gene expression by c-Myc. J Biol Chem 275:21797–800.

9. Ying H, Kimmelman AC, Lyssiotis CA, Hua S, Chu GC, Fletcher-Sananikone E, Locasale JW, Son J, Zhang H, Coloff JL, Yan H, Wang W, Chen S, Viale A, Zheng H, Paik JH, Lim C, Guimaraes AR, Martin ES, Chang J, Hezel AF, Perry SR, Hu J, Gan B, Xiao Y, Asara JM, Weissleder R, Wang YA, Chin L, Cantley LC, DePinho RA. 2012. Oncogenic Kras maintains pancreatic tumors through regulation of anabolic glucose metabolism. Cell 149:656–70.

10. Wu N, Zheng B, Shaywitz A, Dagon Y, Tower C, Bellinger G, Shen CH, Wen J, Asara J, McGraw TE, Kahn BB, Cantley LC. 2013. AMPK-dependent degradation of TXNIP upon energy stress leads to enhanced glucose uptake via GLUT1. Mol Cell 49:1167–75.

11. Waldhart AN, Dykstra H, Peck AS, Boguslawski EA, Madaj ZB, Wen J, Veldkamp K, Hollowell M, Zheng B, Cantley LC, McGraw TE, Wu N. 2017. Phosphorylation of TXNIP by AKT Mediates Acute Influx of Glucose in Response to Insulin. Cell Rep 19:2005–2013.

12. Parikh H, Carlsson E, Chutkow WA, Johansson LE, Storgaard H, Poulsen P, Saxena R, Ladd C, Schulze PC, Mazzini MJ, Jensen CB, Krook A, Bjornholm M, Tornqvist H, Zierath JR, Ridderstrale M, Altshuler D, Lee RT, Vaag A, Groop LC, Mootha VK. 2007. TXNIP regulates peripheral glucose metabolism in humans. PLoS Med 4:e158.

13. Chutkow WA, Patwari P, Yoshioka J, Lee RT. 2008. Thioredoxin-interacting protein (Txnip) is a critical regulator of hepatic glucose production. J Biol Chem 283:2397–406.

14. Hui ST, Andres AM, Miller AK, Spann NJ, Potter DW, Post NM, Chen AZ, Sachithanantham S, Jung DY, Kim JK, Davis RA. 2008. Txnip balances metabolic and growth signaling via PTEN disulfide reduction. Proc Natl Acad Sci U S A 105:3921–6.

15. Stoltzman CA, Peterson CW, Breen KT, Muoio DM, Billin AN, Ayer DE. 2008. Glucose sensing by MondoA:Mlx complexes: a role for hexokinases and direct regulation of thioredoxin-interacting protein expression. Proc Natl Acad Sci U S A 105:6912–7.

16. Chutkow WA, Birkenfeld AL, Brown JD, Lee HY, Frederick DW, Yoshioka J, Patwari P, Kursawe R, Cushman SW, Plutzky J, Shulman GI, Samuel VT, Lee RT. 2010. Deletion of the alpha-arrestin protein Txnip in mice promotes adiposity and adipogenesis while preserving insulin sensitivity. Diabetes 59:1424–34.

17. DeBalsi KL, Wong KE, Koves TR, Slentz DH, Seiler SE, Wittmann AH, Ilkayeva OR, Stevens RD, Perry CG, Lark DS, Hui ST, Szweda L, Neufer PD, Muoio DM. 2014. Targeted metabolomics connects thioredoxin-interacting protein (TXNIP) to mitochondrial fuel selection and regulation of specific oxidoreductase enzymes in skeletal muscle. J Biol Chem 289:8106–20.

18. Saxena G, Chen J, Shalev A. 2010. Intracellular shuttling and mitochondrial function of thioredoxin-interacting protein. J Biol Chem 285:3997–4005.

19. Jeon JH, Lee KN, Hwang CY, Kwon KS, You KH, Choi I. 2005. Tumor suppressor VDUP1 increases p27(kip1) stability by inhibiting JAB1. Cancer Res 65:4485–9.

20. Shen L, O’Shea JM, Kaadige MR, Cunha S, Wilde BR, Cohen AL, Welm AL, Ayer DE. 2015. Metabolic reprogramming in triple-negative breast cancer through Myc suppression of TXNIP. Proc Natl Acad Sci U S A 112:5425–30.

21. Chen JL, Merl D, Peterson CW, Wu J, Liu PY, Yin H, Muoio DM, Ayer DE, West M, Chi JT. 2010. Lactic acidosis triggers starvation response with paradoxical induction of TXNIP through MondoA. PLoS Genet 6:e1001093.

22. Cadenas C, Franckenstein D, Schmidt M, Gehrmann M, Hermes M, Geppert B, Schormann W, Maccoux LJ, Schug M, Schumann A, Wilhelm C, Freis E, Ickstadt K, Rahnenfuhrer J, Baumbach JI, Sickmann A, Hengstler JG. 2010. Role of thioredoxin reductase 1 and thioredoxin interacting protein in prognosis of breast cancer. Breast Cancer Res 12:R44.

23. Peterson CW, Stoltzman CA, Sighinolfi MP, Han KS, Ayer DE. 2010. Glucose controls nuclear accumulation, promoter binding, and transcriptional activity of the MondoA-Mlx heterodimer. Mol Cell Biol 30:2887–95.

24. O’Shea JM, Ayer DE. 2013. Coordination of nutrient availability and utilization by MAX- and MLX-centered transcription networks. Cold Spring Harb Perspect Med 3:a014258.

25. Billin AN, Eilers AL, Coulter KL, Logan JS, Ayer DE. 2000. MondoA, a novel basic helix-loop-helix-leucine zipper transcriptional activator that constitutes a positive branch of a max-like network. Mol Cell Biol 20:8845–54.

26. Sans CL, Satterwhite DJ, Stoltzman CA, Breen KT, Ayer DE. 2006. MondoA-Mlx heterodimers are candidate sensors of cellular energy status: mitochondrial localization and direct regulation of glycolysis. Mol Cell Biol 26:4863–71.

27. Elgort MG, O’Shea JM, Jiang Y, Ayer DE. 2010. Transcriptional and Translational Downregulation of Thioredoxin Interacting Protein Is Required for Metabolic Reprogramming during G(1). Genes Cancer 1:893–907.

28. Kaadige MR, Yang J, Wilde BR, Ayer DE. 2015. MondoA-Mlx transcriptional activity is limited by mTOR-MondoA interaction. Mol Cell Biol 35:101–10.

29. Zhang P, Wang C, Gao K, Wang D, Mao J, An J, Xu C, Wu D, Yu H, Liu JO, Yu L. 2010. The ubiquitin ligase itch regulates apoptosis by targeting thioredoxin-interacting protein for ubiquitin-dependent degradation. J Biol Chem 285:8869–79.

30. Proud CG. 2007. Signalling to translation: how signal transduction pathways control the protein synthetic machinery. Biochem J 403:217–34.

31. Holland EC, Sonenberg N, Pandolfi PP, Thomas G. 2004. Signaling control of mRNA translation in cancer pathogenesis. Oncogene 23:3138–44.

32. Faller WJ, Jackson TJ, Knight JR, Ridgway RA, Jamieson T, Karim SA, Jones C, Radulescu S, Huels DJ, Myant KB, Dudek KM, Casey HA, Scopelliti A, Cordero JB, Vidal M, Pende M, Ryazanov AG, Sonenberg N, Meyuhas O, Hall MN, Bushell M, Willis AE, Sansom OJ. 2015. mTORC1-mediated translational elongation limits intestinal tumour initiation and growth. Nature 517:497–500.

33. Ingolia NT, Lareau LF, Weissman JS. 2011. Ribosome profiling of mouse embryonic stem cells reveals the complexity and dynamics of mammalian proteomes. Cell 147:789–802.

34. Gingold H, Pilpel Y. 2011. Determinants of translation efficiency and accuracy. Mol Syst Biol 7:481.

35. Frenkel-Morgenstern M, Danon T, Christian T, Igarashi T, Cohen L, Hou YM, Jensen LJ. 2012. Genes adopt non-optimal codon usage to generate cell cycle-dependent oscillations in protein levels. Mol Syst Biol 8:572.

36. Novoa EM, Ribas de Pouplana L. 2012. Speeding with control: codon usage, tRNAs, and ribosomes. Trends Genet 28:574–81.

37. Quax TE, Claassens NJ, Soll D, van der Oost J. 2015. Codon Bias as a Means to Fine-Tune Gene Expression. Mol Cell 59:149–61.

38. Tenson T, Ehrenberg M. 2002. Regulatory nascent peptides in the ribosomal tunnel. Cell 108:591–4.

39. Nakatogawa H, Ito K. 2002. The ribosomal exit tunnel functions as a discriminating gate. Cell 108:629–36.

40. Onouchi H, Nagami Y, Haraguchi Y, Nakamoto M, Nishimura Y, Sakurai R, Nagao N, Kawasaki D, Kadokura Y, Naito S. 2005. Nascent peptide-mediated translation elongation arrest coupled with mRNA degradation in the CGS1 gene of Arabidopsis. Genes Dev 19:1799–810.

41. Onouchi H, Haraguchi Y, Nakamoto M, Kawasaki D, Nagami-Yamashita Y, Murota K, Kezuka-Hosomi A, Chiba Y, Naito S. 2008. Nascent peptide-mediated translation elongation arrest of Arabidopsis thaliana CGS1 mRNA occurs autonomously. Plant Cell Physiol 49:549–56.

42. Law GL, Raney A, Heusner C, Morris DR. 2001. Polyamine regulation of ribosome pausing at the upstream open reading frame of S-adenosylmethionine decarboxylase. J Biol Chem 276:38036–43.

43. Raney A, Law GL, Mize GJ, Morris DR. 2002. Regulated translation termination at the upstream open reading frame in s-adenosylmethionine decarboxylase mRNA. J Biol Chem 277:5988–94.

44. Reynolds K, Zimmer AM, Zimmer A. 1996. Regulation of RAR beta 2 mRNA expression: evidence for an inhibitory peptide encoded in the 5′-untranslated region. J Cell Biol 134:827–35.

45. Parola AL, Kobilka BK. 1994. The peptide product of a 5′ leader cistron in the beta 2 adrenergic receptor mRNA inhibits receptor synthesis. J Biol Chem 269:4497–505.

46. Patwari P, Chutkow WA, Cummings K, Verstraeten VL, Lammerding J, Schreiter ER, Lee RT. 2009. Thioredoxin-independent regulation of metabolism by the alpha-arrestin proteins. J Biol Chem 284:24996–5003.

47. Keshwani MM, Klammt C, von Daake S, Ma Y, Kornev AP, Choe S, Insel PA, Taylor SS. 2012. Cotranslational cis-phosphorylation of the COOH-terminal tail is a key priming step in the maturation of cAMP-dependent protein kinase. Proc Natl Acad Sci U S A 109:E1221–9.

48. Dai N, Christiansen J, Nielsen FC, Avruch J. 2013. mTOR complex 2 phosphorylates IMP1 cotranslationally to promote IGF2 production and the proliferation of mouse embryonic fibroblasts. Genes Dev 27:301–12.

49. Ladror DT, Frey BL, Scalf M, Levenstein ME, Artymiuk JM, Smith LM. 2014. Methylation of yeast ribosomal protein S2 is elevated during stationary phase growth conditions. Biochem Biophys Res Commun 445:535–41.

50. Sauert M, Temmel H, Moll I. 2015. Heterogeneity of the translational machinery: Variations on a common theme. Biochimie 114:39–47.

51. Billin AN, Eilers AL, Queva C, Ayer DE. 1999. Mlx, a novel Max-like BHLHZip protein that interacts with the Max network of transcription factors. J Biol Chem 274:36344–50.

52. Bild AH, Yao G, Chang JT, Wang Q, Potti A, Chasse D, Joshi MB, Harpole D, Lancaster JM, Berchuck A, Olson JA, Jr., Marks JR, Dressman HK, West M, Nevins JR. 2006. Oncogenic pathway signatures in human cancers as a guide to targeted therapies. Nature 439:353–7.

53. Pereira B, Chin SF, Rueda OM, Vollan HK, Provenzano E, Bardwell HA, Pugh M, Jones L, Russell R, Sammut SJ, Tsui DW, Liu B, Dawson SJ, Abraham J, Northen H, Peden JF, Mukherjee A, Turashvili G, Green AR, McKinney S, Oloumi A, Shah S, Rosenfeld N, Murphy L, Bentley DR, Ellis IO, Purushotham A, Pinder SE, Borresen-Dale AL, Earl HM, Pharoah PD, Ross MT, Aparicio S, Caldas C. 2016. The somatic mutation profiles of 2,433 breast cancers refines their genomic and transcriptomic landscapes. Nat Commun 7:11479.

54. Cancer Genome Atlas Research N. 2014. Comprehensive molecular profiling of lung adenocarcinoma. Nature 511:543–50.

55. Cerami E, Gao J, Dogrusoz U, Gross BE, Sumer SO, Aksoy BA, Jacobsen A, Byrne CJ, Heuer ML, Larsson E, Antipin Y, Reva B, Goldberg AP, Sander C, Schultz N. 2012. The cBio cancer genomics portal: an open platform for exploring multidimensional cancer genomics data. Cancer Discov 2:401–4.

56. Gao J, Aksoy BA, Dogrusoz U, Dresdner G, Gross B, Sumer SO, Sun Y, Jacobsen A, Sinha R, Larsson E, Cerami E, Sander C, Schultz N. 2013. Integrative analysis of complex cancer genomics and clinical profiles using the cBioPortal. Sci Signal 6:pl1.

57. Ingolia NT, Brar GA, Rouskin S, McGeachy AM, Weissman JS. 2012. The ribosome profiling strategy for monitoring translation in vivo by deep sequencing of ribosome-protected mRNA fragments. Nat Protoc 7:1534–50.

